# WUSCHEL acts as a rheostat on the auxin pathway to maintain apical stem cells in *Arabidopsis*

**DOI:** 10.1101/468421

**Authors:** Yanfei Ma, Andrej Miotk, Zoran Šutiković, Anna Medzihradszky, Christian Wenzl, Olga Ermakova, Christophe Gaillochet, Joachim Forner, Gözde Utan, Klaus Brackmann, Carlos S. Galvan-Ampudia, Teva Vernoux, Thomas Greb, Jan U. Lohmann

## Abstract

To maintain the balance between long-term stem cell self-renewal and differentiation, dynamic signals need to be translated into spatially precise and temporally stable gene expression states. In the apical plant stem cell system, local accumulation of the small, highly mobile phytohormone auxin triggers differentiation while at the same time, pluripotent stem cells are maintained throughout the entire life-cycle. We find that stem cells are resistant to auxin mediated differentiation, but require low levels of signaling for their maintenance. We demonstrate that the WUSCHEL transcription factor confers this behavior by rheostatically controlling the auxin signaling and response pathway. Finally, we show that WUSCHEL acts via regulation of histone acetylation at target loci, including those with functions in the auxin pathway. Our results reveal an important mechanism that allows cells to differentially translate a potent and highly dynamic developmental signal into stable cell behavior with high spatial precision and temporal robustness.

## INTRODUCTION

The shoot apical meristem (SAM) is a highly dynamic and continuously active stem cell system responsible for the generation of all above ground tissues of plants. The stem cells are located in the central zone and are maintained by a feedback loop consisting of the stem cell promoting WUSCHEL (WUS) homeodomain transcription factor and the restrictive CLAVATA (CLV) pathway^1,2^. WUS protein is produced by a group of niche cells, called organizing center, localized in the deeper tissue layers of the meristem ^3^ and moves to stem cells via plasmodesmata^4,5^. WUS is required for maintaining stem cells and SAMs of *wus* mutants terminate due to stem cell exhaustion after producing a small number of organs^6^. Conversely, mutants in genes of the *CLV* pathway exhibit substantial stem cell over-proliferation, which is strictly dependent on *WUS* activity^1,2^. *CLV3* is the only component of this system that is specifically expressed in stem cells and hence serves as a faithful molecular marker. Stem cells are surrounded by transient amplifying cells, which are competent to undergo differentiation in response to auxin, a small, mobile signaling molecule with diverse and context specific roles in plant development and physiology (reviewed in ref. 7). Auxin sensing is dependent on nuclear receptors including *TRANSPORT INHIBITOR RESPONSE1 (TIR1)*, whose activation triggers the proteolytic degradation of AUX/IAA proteins, such as BODENLOS (BDL). AUX/IAA proteins repress auxin responses by inhibiting the function of activating AUXIN RESPONSE FACTOR (ARF) transcription factors via dimerization^8-10^. Intracellular accumulation of auxin is regulated by active polar transport and in the context of the SAM, the export carrier PINFORMED1 (PIN1) determines the sites of lateral organ initiation and thus differentiation^11,12^. In addition to promoting organ initiation, auxin influences stem cell proliferation by interacting with the signaling cascade of another classical phytohormone, cytokinin, and allows lateral organs to communicate with the center of the meristem^13-15^. Here we ask how long-term stem cell fate is robustly maintained within a tissue environment that is subject to such a highly dynamic signaling system geared towards differentiation.

## RESULTS

### Role of auxin signaling for apical stem cell fate

To analyze auxin distribution and response with cellular resolution across the homeostatic apical stem cell system of *Arabidopsis*, we mapped auxin signaling behavior using the genetically encoded markers R2D2 and DR5v2^16^. R2D2 is based on a fusion of the auxin-dependent degradation domain II of an Aux/IAA protein to Venus fluorescent protein, and uses a mutated, non-degradable domain II linked to tdTomato as an internal control^16^. Hence, R2D2 signal is dictated by the levels of auxin as well as the endogenous receptors and represents a proxy for the auxin signaling input for every cell. Following multispectral live-cell image acquisition in plants carrying R2D2, we used computational analysis of the green to red ratio to determine the cellular auxin input status. We found that auxin is present and sensed fairly uniformly across the SAM including the central stem cell domain, with local minima only detected at organ boundaries (Fig. 1a, b and refs. 17,18). In contrast, DR5v2, a reporter for auxin signaling output based on a synthetic promoter containing repeats of ARF DNA binding motifs, was strongly activated non-uniformly in wedge shaped zones of differentiation competent cells, but only weakly expressed the center of the SAM (Fig. 1d; ref. 17). To spatially correlate cellular auxin output status with stem cell fate, we combined the DR5v2 reporter with a *pCLV3:mCherry-NLS* marker in a single transgenic line. Computational analysis of the DR5v2 and *pCLV3* signals revealed that the auxin response minimum invariantly coincided with the center of the stem cell domain (Fig. 1c-f).

**Figure 1:**
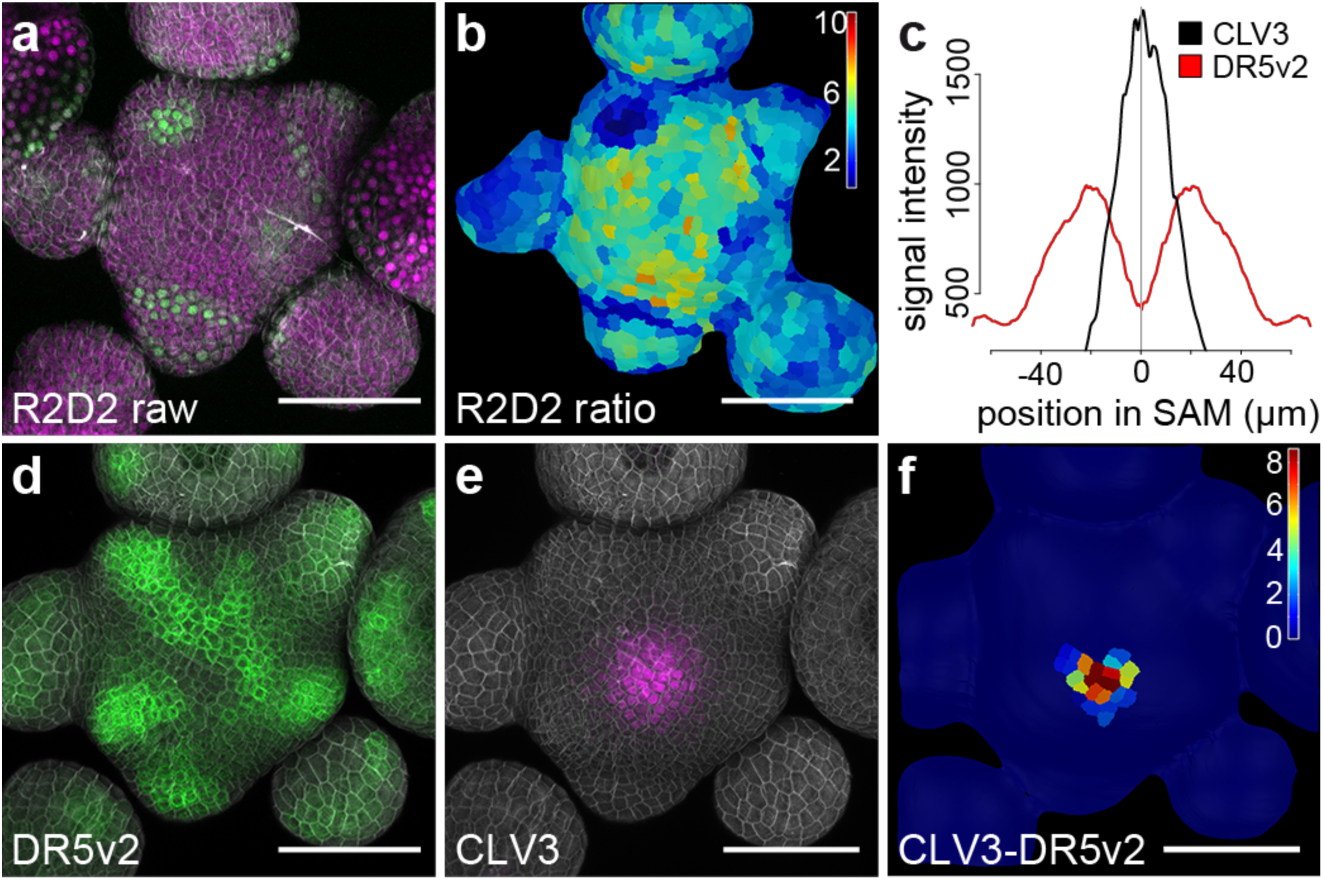
Auxin output minimum correlates with apical stem cells. **a**) Confocal readout from R2D2 auxin input sensor. **b**) Ratiometric representation of R2D2 activity in the epidermal cell layer (L1). **c**) Quantification of averaged *pDR5v2:ER-eYFP-HDEL* and *pCLV3:mCherry-NLS* distribution (n×5). **d**) Confocal readout from *pDR5v2:ER-eYFP-HDEL* auxin output reporter. **e**) *pCLV3:mCherry-NLS* stem cell marker in the same SAM. **f**) Computational subtraction of L1 signals shown in (d) and (e). Relative signal intensity is shown in arbitrary units. Scale bars: 50 *µ*m.

To test if the auxin output minimum is functionally connected to stem cell identity, we interfered with their maintenance. To this end, we experimentally induced symplastic isolation through callose deposition at plasmodesmata of stem cells, which we had shown earlier to induce their differentiation^5,19^. Following DR5v2 signal over time, we observed activation of auxin signaling output in the central zone domain after 36 hours of callose synthase (iCalSm) expression. In addition, cell expansion, a hallmark of plant cell differentiation, became obvious after 72 hours (Fig. 2a-d). All plants that exhibited stem cell loss following to iCalSm activation showed this pattern, which also led to a significant increase in DR5v2 signal intensity over time, in contrast to controls that did not respond (Fig. 2e-g; Supplementary Fig. 1).

**Fig. 2:**
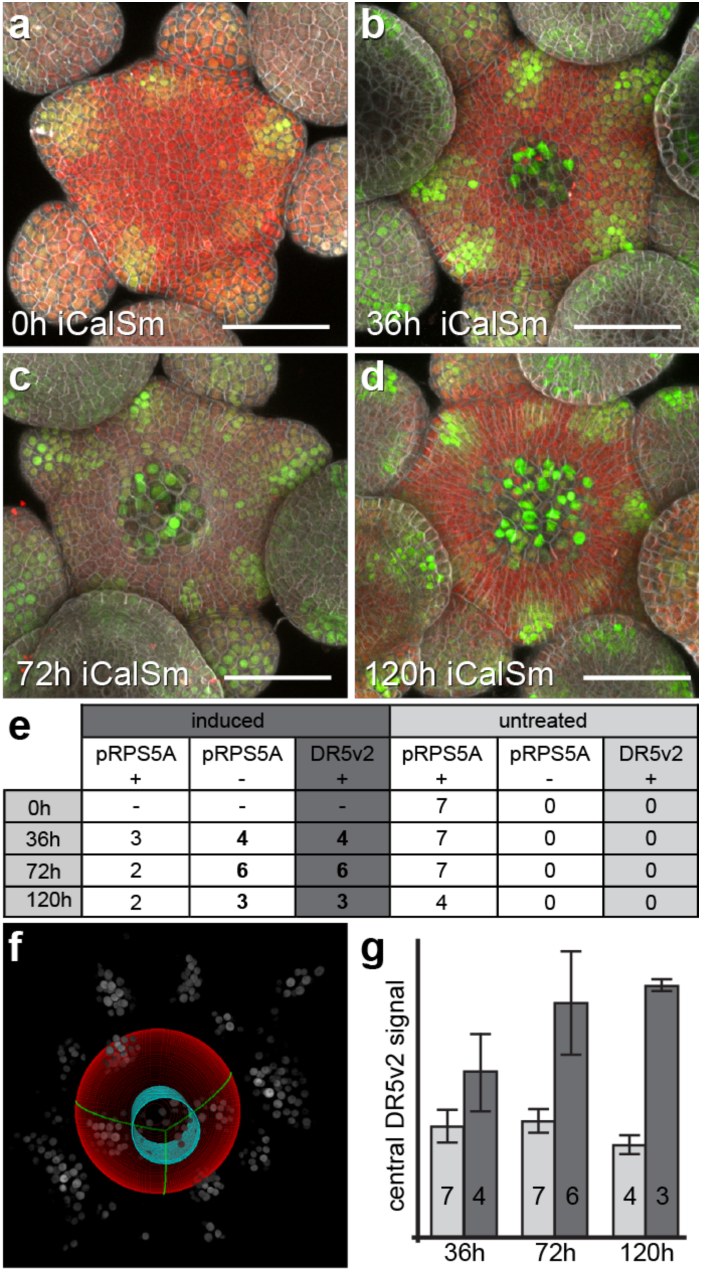
The central auxin signaling minimum is dependent on stem cell fate. **a-d**) *pDR5v2:3xVENUS-NLS* activity after induction of iCalSm. Stem cell differentiation is marked by loss of *pRPS5a:NLS-tdTomato.* **e)** Quantification of DR5v2 response to induction of iCalSm at the per plant level. Number of plants scored for loss of *RPS5a* promoter activity from stem cells and DR5v2 expression are shown. Stem cell loss and associated DR5v2 activation exclusively occurred in induced plants. All plants with stem cell loss as shown by reduced *pRPS5a* activity expressed DR5v2. *pRPS5a* + denotes plants with uncompromised *pRPS5a* promoter activity in stem cells. *pRPS5a* - denotes plants with reduced *pRPS5a* promoter activity in stem cells. DR5v2 + denotes plants with DR5v2 activity in stem cells. **f)** Computational sphere fitting and identification of the central zone for fluorescence signal quantification. **g)** Quantification of DR5v2 signal intensity in the central zone across the experimental cohort described in (e). Light grey bars represent uninduced controls, dark grey bars represent plants induced with 1% ethanol. Numbers of analyzed SAMs are indicated. See also Supplementary Figure 1.

Thus, stem cell fate and the auxin response minimum appeared to be functionally connected, leading us to hypothesize that manipulation of auxin signaling in the central zone should affect stem cell behavior. To test this directly, we designed a transgene to suppress auxin signaling output specifically in stem cells. Therefore, we fused the dominant auxin signaling output inhibitor *BDL-D* (IAA12) with the glucocorticoid receptor tag. The activity of the resulting fusion protein could be induced by dexamethasone (DEX) treatment, which allowed the translocation of BDL-D-GR from the cytoplasm to the nucleus, its native cellular compartment^20^. In line with our expectations, we found that inducing *pCLV3:BDL-D-GR* led to an expansion of the DR5v2 minimum in the center of the SAM reflecting the inhibitory activity of BDL-D on ARF transcription factors (Fig. 3a, b). Surprisingly, long term induction of BDL-D-GR or stem cell specific expression of *BDL-D* without the GR tag caused meristem termination in about half of the seedlings (n×90; Fig. 3f, g), demonstrating that stem cells require active auxin signaling for their maintenance. In contrast, expression of a potent positive signaling component, the auxin response factor *ARF5/MONOPTEROS (MP)*, or its constitutively active form *MPΔ*, which engages the auxin pathway independently of signal perception^21^, did not cause relevant reduction in meristem size (Fig. 3c-e, h, j and ref. 15). When expressed throughout the entire SAM by the HMG promoter (Supplementary Fig. 2a, b), *MPΔ* stimulated ectopic organ initiation specifically in the peripheral zone (Fig. 3i), demonstrating that resistance to auxin was not a general feature of the meristem, but limited to stem cells. Importantly, the DR5v2 reporter, which senses auxin output by providing binding sites for ARF transcription factors, was activated in stem cells of plants expressing *MP* and *MPΔ* (6/8 independent T1 lines) (Fig. 3c-e and Supplementary Fig. 2c-k), suggesting that the resistance to auxin occurs, at least in part, downstream of ARF activity. Taken together, these experiments demonstrated that auxin signaling is locally gated to permit a low instructive output level, while at the same time protecting stem cells from the differentiation inducing effects of the phytohormone at high signaling levels.

**Fig. 3:**
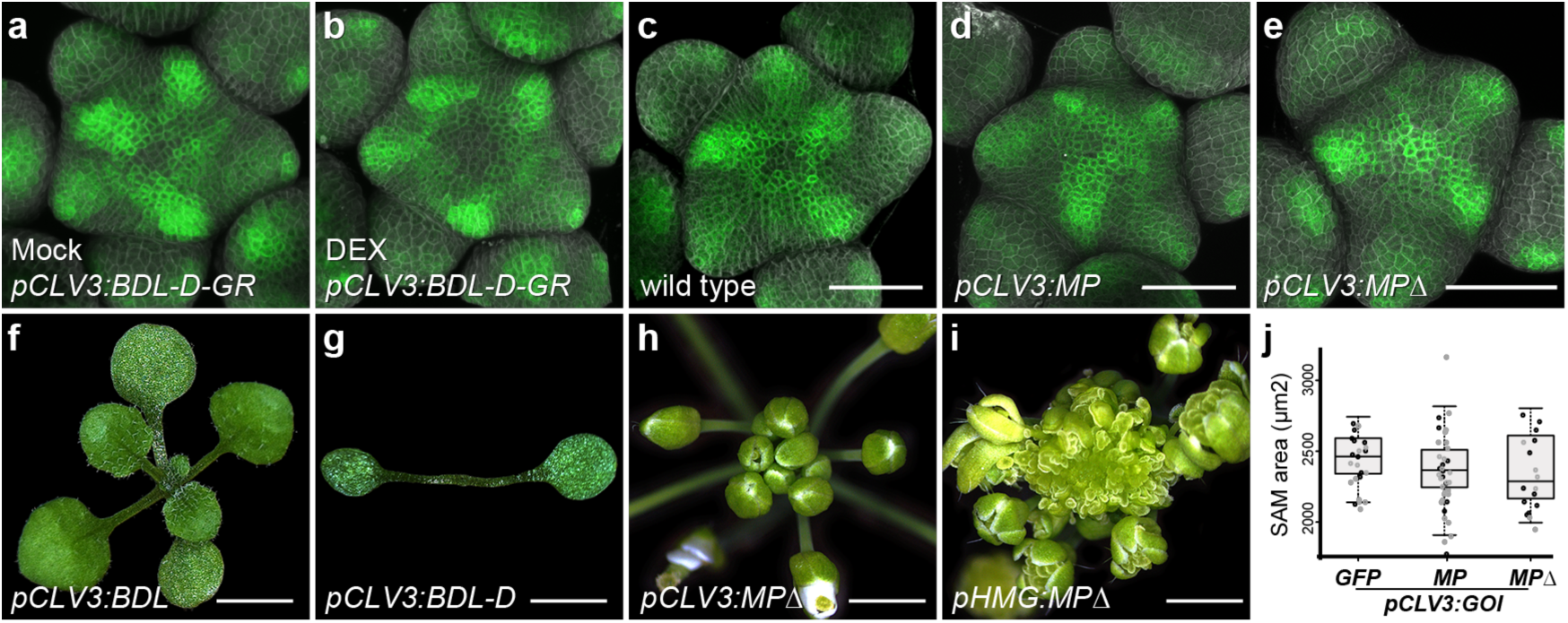
Stem cells require auxin signaling, but are resistant to overactivation of the pathway. **a-e**) *pDR5v2:ER-eYFP-HDEL* activity in plants harboring *pCLV3:BDL-D-GR* after 24h of mock treatment (A), *pCLV3:BDL-D-GR* after 24h of DEX treatment (B), wild type (C), *pCLV3:MP* (D) or *pCLV3:MPΔ* (E). **f-i**) Representative phenotypes of lines expressing *pCLV3:BDL* (F), *pCLV3:BDL-D* (G), *pCLV3:MPΔ* (H), or *pHMG:MPΔ* (I). **j)** SAM size quantifications for plants carrying *pCLV3:GFP, pCLV3:MP*, or *pCLV3:MPΔ* in two independent T1 populations. All scale bars 50 *µ*m, except F) and G) 3,5 mm; H) and I) 2mm.

### WUSCHEL controls auxin signaling output in stem cells

Since suppressing auxin signaling output in stem cell caused SAM arrest and a phenotype highly similar to *wus* mutants (Fig. 3f, g), we tested the contribution of *WUS* to controlling auxin responses in diverse genetic backgrounds. The *WUS* expression domain is massively enlarged in *clv* mutants^1,2^, which causes stem cell over-proliferation phenotypes, and therefore SAMs from these plants provide an ideal background to elucidate the functional connection of WUS and auxin. Consequently, we analyzed auxin output in *clv3* meristems and found the DR5v2 minimum expanded in line with the overaccumulation of WUS, however some weak signal remained throughout the SAM (Fig. 4a, b). To test whether auxin signaling is required for stem cell over-proliferation in *clv3* mutants, we locally blocked auxin output by our *pCLV3:BDL-D* transgene and observed stem cell termination phenotypes in almost all seedlings (n×30; Fig. 4c). This result suggested that also in fasciated SAMs of *clv3* mutants, ectopic WUS is sufficient to reduce auxin signaling, while at the same time permitting basal output levels. To test the short term effect of enhancing WUS levels without the indirect effects of the *clv3* phenotype, we created plants that carry a *pUBI10:mCherry-GR-linker-WUS (WUS-GR)* transgene which allowed for experimental induction of ubiquitous WUS activity. After 24 h of DEX treatment the central auxin signaling minimum as well as the *CLV3* domain expanded (Fig. 4d-f; Supplementary Fig. 3a-f), suggesting that WUS is indeed sufficient to reduce signaling output in the center of the SAM, but is unable to override active auxin responses at the periphery. To test whether *WUS* is also required to protect stem cells from high signaling levels, which lead to differentiation, we developed a genetic system that allowed us to inducibly degrade WUS protein in stem cells. To this end, we adapted deGradFP technology^22^ and combined switchable stem cell specific expression of an anti-GFP nanobody with a *pWUS:WUS-linker-GFP wus* rescue line^5^. After 24h induction of nanobody expression, WUS-linker-GFP signal was substantially reduced in stem cells of the epidermis and subepidermis (Fig. 4g-h) and after five days we observed shoot termination (Fig. 4i). Combining this *wus/pWUS:WUS-linker-GFP/pCLV3:AlcR/pAlcA:NSlmb-vhhGFP4* line with the DR5v2 marker showed that after 24h of WUS depletion, cells in center of the SAM had become responsive to auxin whereas they remained insensitive in mock treated controls (Fig. 4j-l). We made similar observations in plants carrying DR5v2 and the weak *wus-7* allele, which were able to maintain a functional SAM for some time and only terminated stochastically. In these lines, DR5v2 activity fluctuated substantially and was frequently observed in the central zone (Fig. 4l and Supplementary Fig. 4). Taken together, these results demonstrated that WUS is required to rheostatically maintain stem cells in a state of low auxin signaling.

**Fig. 4:**
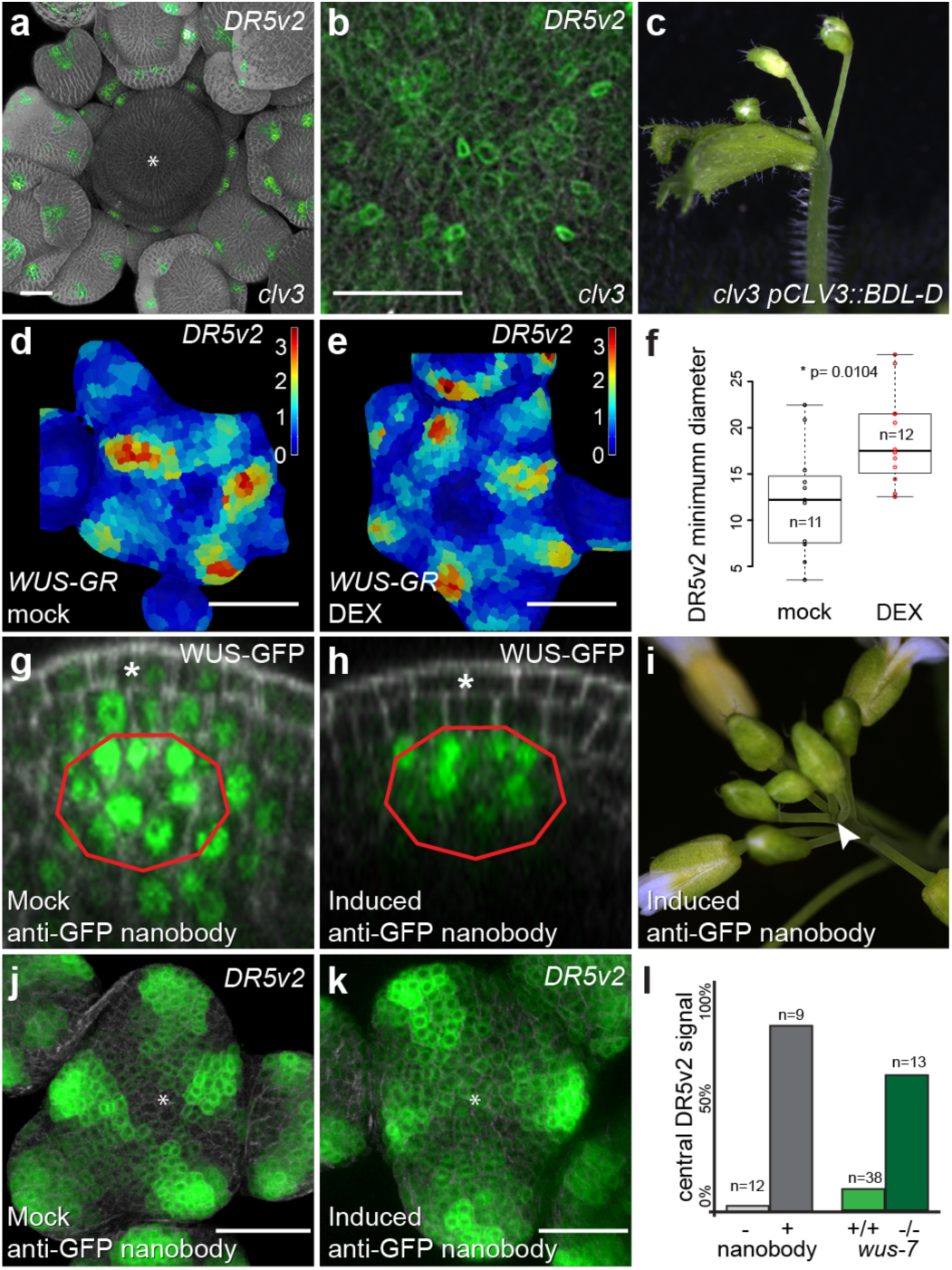
WUSCHEL maintains low auxin signaling output in stem cells. **a**) *pDR5v2:ER-mCherry-HDEL* activity in SAM of *clv3* mutant. Asterisk marks center of SAM. **b**) Zoom into central SAM area of *clv3* mutants reveals basal *pDR5v2* activity. **c**) SAM arrest caused by *pCLV3:BDL-D* expression in *clv3*. **d, e**) Representative *pDR5v2:ER-mCherry-HDEL* signals after 24h of mock treatment (D) or inducible ectopic activation of *WUS-GR* activity (E). **f**) Quantification of central DR5v2 signal minimum following ectopic WUS activation. **g, h**) Representative images of a *pWUS:WUS-linker-GFP* rescue line expressing the anti GFP nanobody under the control of *pCLV3:AlcR* (*wus/pWUS:WUS-linker-GFP/pCLV3:AlcR/pAlcA:NSlmb-vhhGFP4)*. **g)** WUS-linker-GFP signal after 24h of mock treatment. **h)** WUS-linker-GFP signal after 24h of WUS depletion. **i)** Shoot termination observed five days after WUS depletion. Red lines mark *WUS* mRNA expressing cells of the organizing centre; asterisk denote epidermal stem cells. **j, k**) Representative *pDR5v2:ER-mCherry-HDEL* signals after 24h of mock treatment (D) or depletion of WUS protein from stem cells. **l)** Quantification of DR5v2 presence in the central zone following WUS depletion or in weak *wus-7* mutants. Scale bars: 50 *µ*m.

### Mechanisms of auxin pathway gating

To address how WUS is able to control the output of the auxin pathway, we went on to define direct target genes combining new ChIP-seq and RNA-seq experiments using seedlings of our *WUS-GR* line. Leveraging the uniform expression and high inducibility of our transgene, as well as the high affinity of RFP-trap single chain antibodies to the mCherry tag used for our ChIP protocol, we were able to identify 6740 genomic regions bound by WUS. This compared to 136 regions we had previously identified by ChIP-chip^23^. Previously identified direct targets, such as *ARR7, CLV1, KAN1, KAN2 AS2* and *YAB3* (refs. 23-25) were also picked up in our new datasets. Interestingly, WUS binding was almost exclusively found in regions of open chromatin^26^ and among the WUS targets we found the gene ontology term “response to auxin” to be most highly enriched within the developmental category (Supplementary Table 1). Importantly, WUS appeared to control auxin signaling output at all relevant levels, since it was able to bind to the promoters or regulate the expression of a large number of genes involved in auxin transport, auxin perception, auxin signal transduction, as well as auxin response, which occurs downstream of ARF transcription factors (Fig. 5a and Supplementary Table 2). Since WUS can act as transcriptional activator or repressor dependent on the regulatory environment^27,28^ and our profiling results were based on ectopic expression of WUS in non-stem cells, we were unable to predict how the expression of individual targets would be affected *in vivo*. However, it has been reported that in the SAM, WUS mainly acts as a transcriptional repressor^23-25,27^ and consistently, many auxin signaling components are expressed at high levels only in the periphery of the SAM and exhibit low RNA accumulation in the cells that are positive for WUS protein^17^. To test if WUS is required for this pattern, we analyzed the response of *MP* and *TIR1* mRNA accumulation to variations in *WUS* expression. To circumvent morphological defects of stable *wus* mutants, we again made use of our deGradFP line to analyze expression of *MP* after loss of WUS protein activity, but prior to changes in SAM morphology. After 24 h of WUS depletion, *MP* mRNA expression had extended from the periphery into the central zone (Fig. 5b, c; Supplementary Fig. 5), demonstrating that WUS is indeed required for *MP* repression in stem cells. Conversely, ectopic activation of WUS revealed that it is also sufficient to reduce, but not shut down *MP* and *TIR1* transcription even in the periphery of the SAM (Fig. 5d-e, Supplementary Fig. 3g, h).

**Table 1:** WUS targets functionally tested by stem cell specific expression. Expression domains in the SAM are based on refs. 17,40,41.

| AGI | Name | Responsive to auxin | Expression PZ>CZ | Promoter bound by WUS | Responsive to WUS |
| --- | --- | --- | --- | --- | --- |
| AT3G62980 | *TIR1* | x | x | x | x |
| AT2G33860 | *ARF3* | x | x | x | x |
| AT5G60450 | *ARF4* | x | x | x | x |
| AT1G19850 | *ARF5 (MP)* | x | x | x | x |
| AT2G22670 | *IAA8* | x | x | - | x |
| AT5G65670 | *IAA9* | x | x | x | x |
| AT1G04550 | *IAA12 (BDL)* | x | x | - | - |
| AT5G60200 | *TMO6* | x | x | x | x |
| AT1G74500 | *TMO7* | x | - | - | - |
| AT3G25710 | *TMO5* | x | - | - | x |
| AT4G23750 | *TMO3* | x | - | x | x |
| AT1G68510 | *LBD42* | - | - | x | - |
| AT3G49940 | *LBD38* | - | - | x | x |
| AT3G58190 | *LBD29* | x | - | - | - |
| AT3G11280 |  | x | x | x | x |
| AT3G28910 | *MYB30* | x | x | x | x |
| AT5G58900 |  | x | x | x | - |

**Fig. 5:**
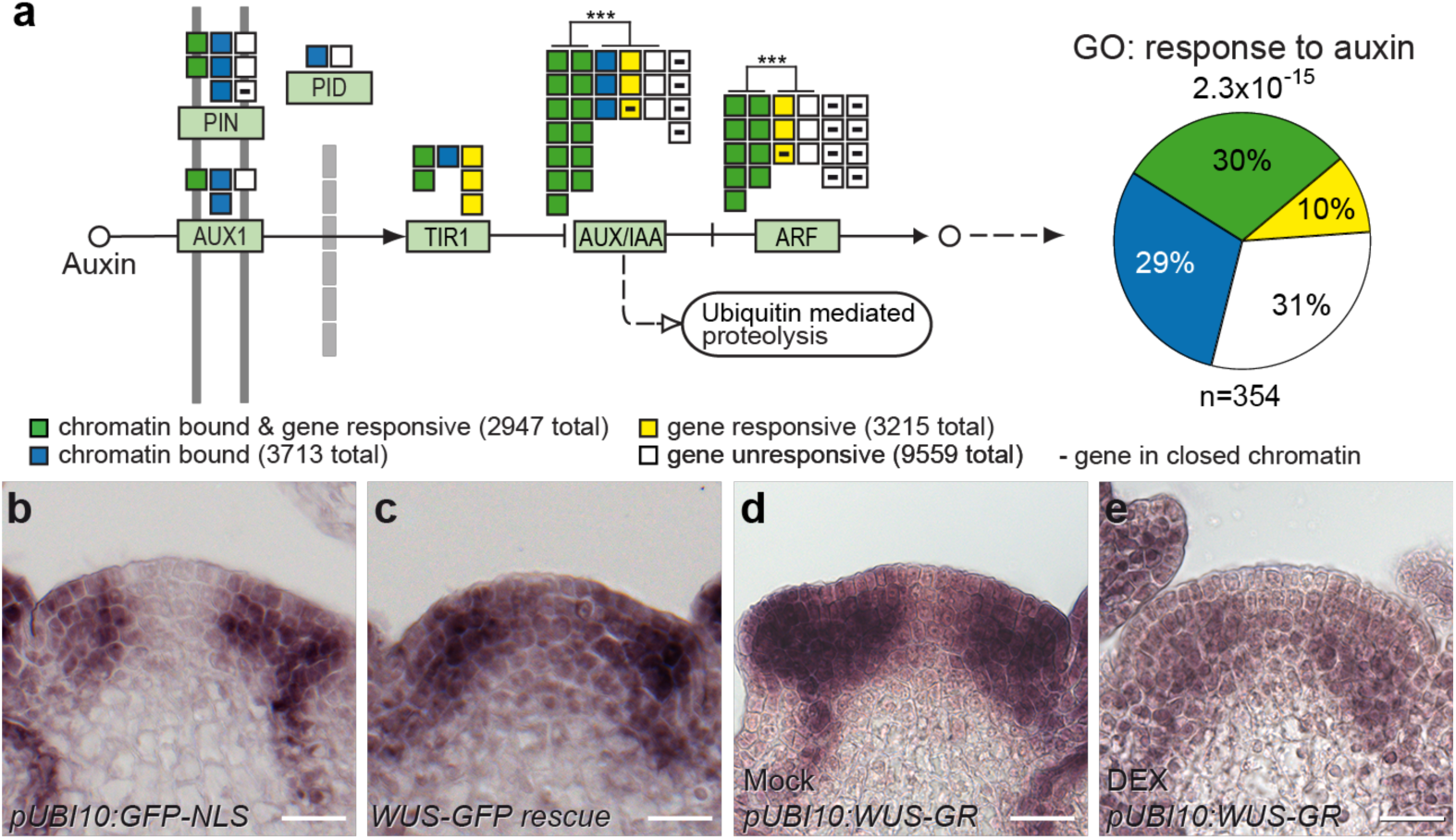
Pathway level control underlies WUSCHEL mediated gating of auxin signaling. **a**) WUS globally affects the auxin pathway, including transport, perception, signal transduction, as well as transcriptional response. Across the entire pathway bound and responsive genes are overrepresented (p-value 9.9*10^−10^). Within gene family tests are shown. *** p-value by Fisher exact test < 10^−4^. **b, c**) *MP* RNA accumulation 24 hours post anti-GFP nanobody induction in a *pUBI10:GFP-NLS* control line (B) and the *pWUS:WUS-linker-GFP wus* rescue background (C). **d, e**) Response of *MP* mRNA to ectopic activation of WUS-GR. *MP* RNA after 24h of mock (D) or DEX treatment (E). Scale bars: 20*µ*m.

To elucidate the molecular mechanisms responsible for the observed rheostatic activity, we asked whether chromatin structure may be changed in response to WUS. WUS physically interacts with TOPLESS (TPL)^29,30^, a member of the GROUCHO/Tup1 family of transcriptional co-repressors. These adaptor proteins mediate interaction with HISTONE DEACETYLASES (HDACs, reviewed in 31), which in turn act to reduce transcriptional activity of chromatin regions via promoting the removal of acetyl modifications from histone tails^32^. To test whether regulation of chromatin modification is involved in translating WUS activity into the observed reduction of transcriptional activity of target genes we quantified histone acetylation on H3K9/K14 and methylation on H3K27. After 2 h of induction of our *WUS-GR* line, we observed a significant change in the genome wide histone acetylation patterns, which were spatially correlated with WUS chromatin binding events (2939 out of 6740 WUS bound chromatin regions showed acetylation changes), while histone methylation patterns were largely unaffected (634 out of 6740 WUS bound chromatin regions showed methylation changes) (Fig. 6a). WUS binding events clustered in the proximal promoter regions, while chromatin regions whose acetylation levels were changed after WUS activation were mainly found around the transcriptional start sites and 5′UTRs of genes (Fig. 6b). Zooming in on the 1656 directly repressed WUS targets, we found that 587 of them also showed histone de-acetylation. For the vast majority of these loci the observed reduction was fairly subtle, suggesting that mild de-acetylation may be the mechanism that allows WUS to reduce, but not shut off transcription of target genes. To test whether the observed changes in chromatin state of direct WUS targets also translate to variation in gene expression, we induced WUS activity in the absence or presence of Trichostatin A (TSA), a potent inhibitor of class I and II HDACs^33^, and recorded the transcriptional response. Principle Component Analysis (PCA) not only showed that both WUS activation and TSA contributed to gene expression variance, but that there was a clear interaction of their activities. Strikingly, roughly 40% of gene expression variance caused by WUS activation was suppressed by TSA treatment (Fig. 6c). Consistently, from the 1656 directly repressed genes, 938 were no longer responsive to WUS-GR induction when TSA was present and roughly a third of them showed significant reduction in H3K9/K14 acetylation levels (Fig. 6d). These results underlined the relevance of histone de-acetylation for the genome-wide functional output of WUS and prompted us to investigate whether this mechanism is relevant for controlling auxin responses in the SAM. Therefore, we analyzed DR5v2 reporter activity after TSA and/or auxin treatment and found that auxin was insufficient to trigger a transcriptional response in stem cells, likely due to the presence of functional WUS (Fig. 6e). In contrast, inactivation of HDACs and consequently WUS-mediated transcriptional repression by TSA treatment, led to low but consistent DR5v2 signal in the center of the meristem (Fig. 6f). Finally, combining a reduction in WUS function by TSA with stimulation of the auxin pathway caused a substantial DR5v2 response in stem cells (Fig. 6g). Taken together, these results showed that WUS binds to and reduces transcription of the majority of genes involved in auxin signaling and response via de-acetylation of histones and thus is able to rheostatically maintain pathway activity in stem cells at a basal level.

**Fig. 6:**
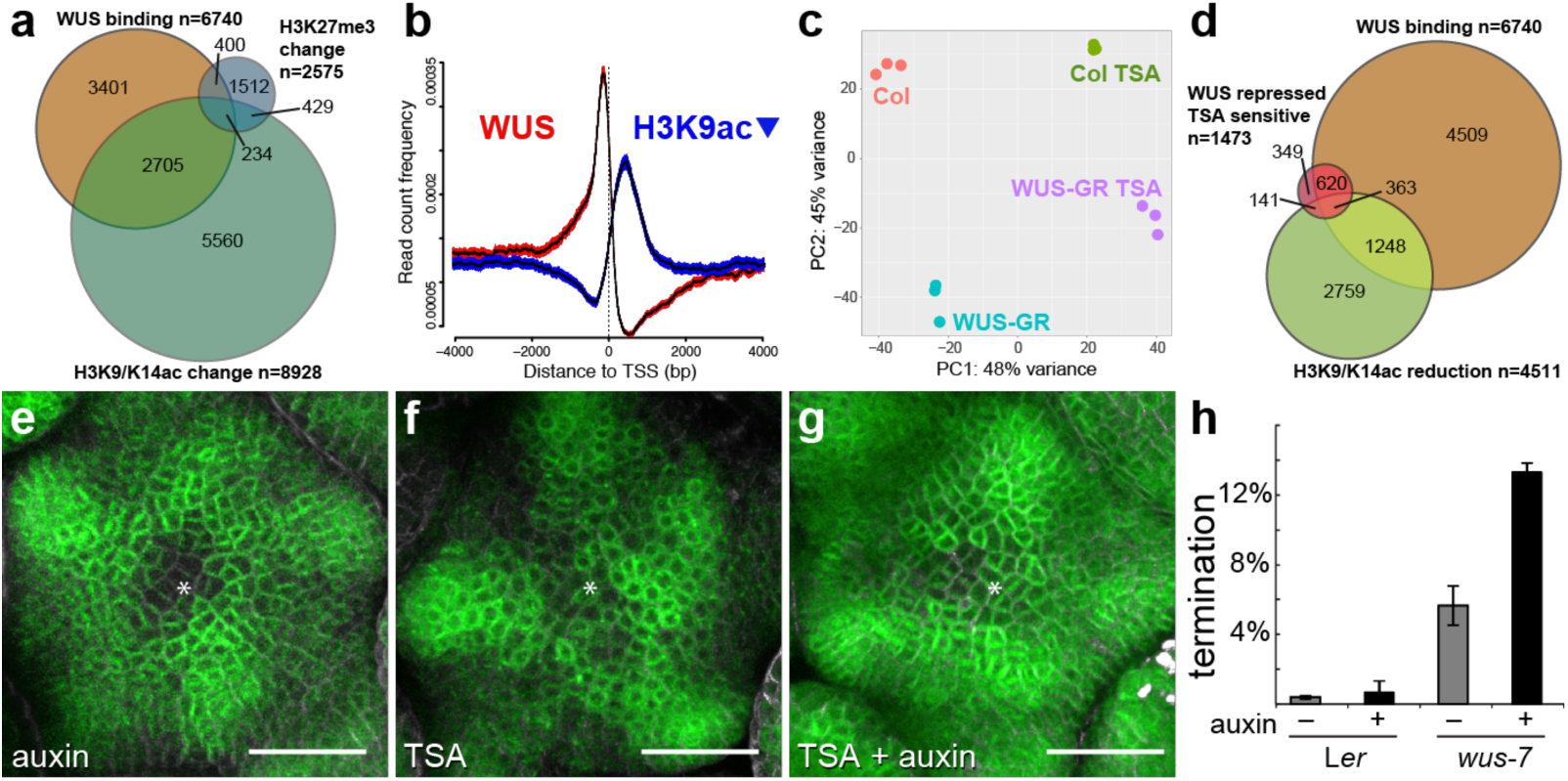
WUS acts by regulating the histone acetylation status of target loci. **a)** Venn diagram showing the overlap between WUS binding regions (orange), and loci with significant changes in H3K9K14ac (green) or H3K27me3 (blue) status. **b)** Spatial correlation between WUS chromatin binding events (red) and regions with reduced histone acetylation (blue) 0.95 confidence intervals are shown. **c)** PCA showing the global transcriptional response to WUS-GR activation in the presence or absence of TSA. TSA treatment suppressed almost 50% of gene expression variance caused by activation of WUS-GR. **d)** Venn diagram showing the overlap between WUS binding regions (orange), and loci with significant reduction in H3K9K14ac (light green) and genes whose expression was reduced by WUS in a TSA sensitive manner (red). **e-g**) Representative images of *pDR5v2:ER-mCherry-HDEL* activity in response to HDAC inhibition. **e)** Auxin treated SAM; **f)** TSA treated SAM; **g)** TSA and auxin treated SAM. Asterisk denote center of the SAM. **h)** Quantification of terminated seedlings grown on auxin plates (10 *µ*M IAA; n > 200 for each genotype and treatment). Genotyping revealed that all arrested plants were homozygous for *wus-7*. Scale bars: 30*µ*m.

### Pathway wide control provides robustness to apical stem cell fate

We next wondered what the functional relevance of the observed pathway wide regulatory interaction might be. Therefore, we tested the capacity of WUS targets with auxin signaling or response functions to interfere with stem cell activity. Based on their highly localized expression at the periphery of the SAM^17^, we selected the signaling components *ARF3, ARF4, ARF5 (MP), IAA8, IAA9*, and *IAA12* (*BDL)* as well as the TIR1 receptor along with transcription factors of the auxin response category including *TARGET OF MONOPTEROS* (*TMO*) and *LATERAL ORGAN BOUNDARIES* (*LOB*) genes that have established roles in other developmental contexts^34^. Neither of the 17 factors tested caused meristem phenotypes when expressed in stem cells (Fig. 2 and Table 1), highlighting the robustness of stem cell fate in the presence of WUS on the one hand and the activity of auxin signaling in these cells on the other hand. This conclusion is based on two observations: 1. The auxin sensitive native version of BDL was unable to terminate the SAM in contrast to the auxin insensitive BDL-D version (Fig. 3f, g). 2. *pCLV3:MP* plants showed enhanced DR5v2 activity in stem cells (Fig. 3c, d) demonstrating that ARF activity is indeed limiting for transcriptional output in wild-type. However, the transcriptional output registered by the DR5v2 reporter was not translated into an auxin response, since WUS limited the expression of a large fraction of the required downstream genes (Fig. 5a; Supplementary Table 2). Thus, WUS seems to act both up- and downstream of the key ARF transcription factors.

Since we had found that stem cell specific expression of individual auxin signaling components was not sufficient to interfere with stem cell fate, we wanted to test whether reducing *WUS* function would sensitize stem cells to activation of the entire pathway. To this end, we grew plants segregating for *wus-7* on plates supplemented with auxin. Eleven days after germination, we observed twice as many terminated *wus-7* mutant seedlings on auxin plates compared to control plates, whereas wild-type seedlings were unaffected (Fig. 6h). Thus, reducing *WUS* function allowed activation of auxin responses under conditions that were tolerated in wild type. Taken together, the activation of individual pathway components was insufficient to override the protective effect of WUS, however compromising the master regulator itself rendered stem cells vulnerable to even mild perturbations in auxin signaling.

## Discussion

In conclusion, our results show that WUS restricts auxin signaling in apical stem cells by pathway-wide transcriptional control, while at the same time allowing instructive low levels of signaling output. This rheostatic activity may be based on selective transcriptional repression/activation of a subset of signaling and response components that render the pathway unresponsive to high input levels. Alternatively, WUS may be able to reduce expression of targets rather than to shut off their activity completely, leaving sufficient capacity for low level signaling only. In support of the latter hypothesis, we demonstrate that WUS acts via de-acetylation of histones and that interfering with HDAC activity triggers auxin responses in stem cells. However, there is evidence supporting both scenarios^23,25,27,28^ and likely both mechanisms work hand in hand dependent on the regulatory environment of the individual cell. Thus, a definitive answer will require inducible WUS loss of function approaches in stem cells coupled with time-resolved whole genome transcript profiling at the single cell level. Importantly, in addition to its effects on auxin signaling, WUS enhances cytokinin responses via the repression of negative feedback regulators^24^. This interaction can be overridden by expression of constitutively active versions of these negative feedback components^24^, and similarly we find here that dominant negative auxin regulators lead to SAM arrest. In contrast, wild-type or constitutively active auxin signaling elements do not lead to SAM defects, suggesting that WUS acts primarily to limit auxin responses. Thus, by acting on both pathways by direct reduction of target gene expression, WUS protects stem cells from auxin mediated differentiation, while at the same time enhancing cytokinin output, which may primarily serve to sustain *WUS* expression^35,36^. Auxin and cytokinin signaling are directly coupled also in other stem cell systems and balancing their outputs is key to maintaining functional plant stem cell niches^15,37^. Given the dynamic and self-organizing nature of the auxin system in the SAM^38^, the independent spatial input provided by WUS appears to be required to bar differentiation competence from the center of the SAM, while at the same time still allowing to sense this important signal. In light of the findings that PIN1 mediated auxin flux in the SAM may be directed towards the center^39^, it is tempting to speculate that auxin could serve as a positional signal not only for organ initiation, but also for stem cells.

## Author Contributions

A. Me. performed in situ hybridizations, C.W. carried out imaging and analyses, J.F. established the WUS-GR line, G.U. and A. M. performed RNA-seq, O.E. performed bioinformatic analyses, K.B. and T.G. established the *pDR5v2:ER-EYFP-HDEL:tAt4g24550* line, C.G. made the *pCLV3:mCherry-NLS:tCLV3* construct, Z.Š., A.M and Y.M. performed all other experiments. C.G.-A. and T.V. designed the TSA treatment of the SAM, Y.M., Z.Š., A.M. and J.U.L. designed all other experiments and wrote the paper with input from all other authors. Sequencing data is available under GEO accession GSE122611.

## Acknowledgments

We thank Dolf Weijers for sharing R2D2 and DR5v2 resources before publication. This work was supported by the DFG through grants SFB1101 and SFB873 to JUL and TG and by HFSP Grant RPG0054-2013 and ANR-12-BSV6-0005 grant to T.V. Computational analyses have been carried out on heiCLOUD provided by Heidelberg University Computing Centre.

## Supplementary Data

### Supplementary Figures 1-5

**Supplementary Figure 1:**
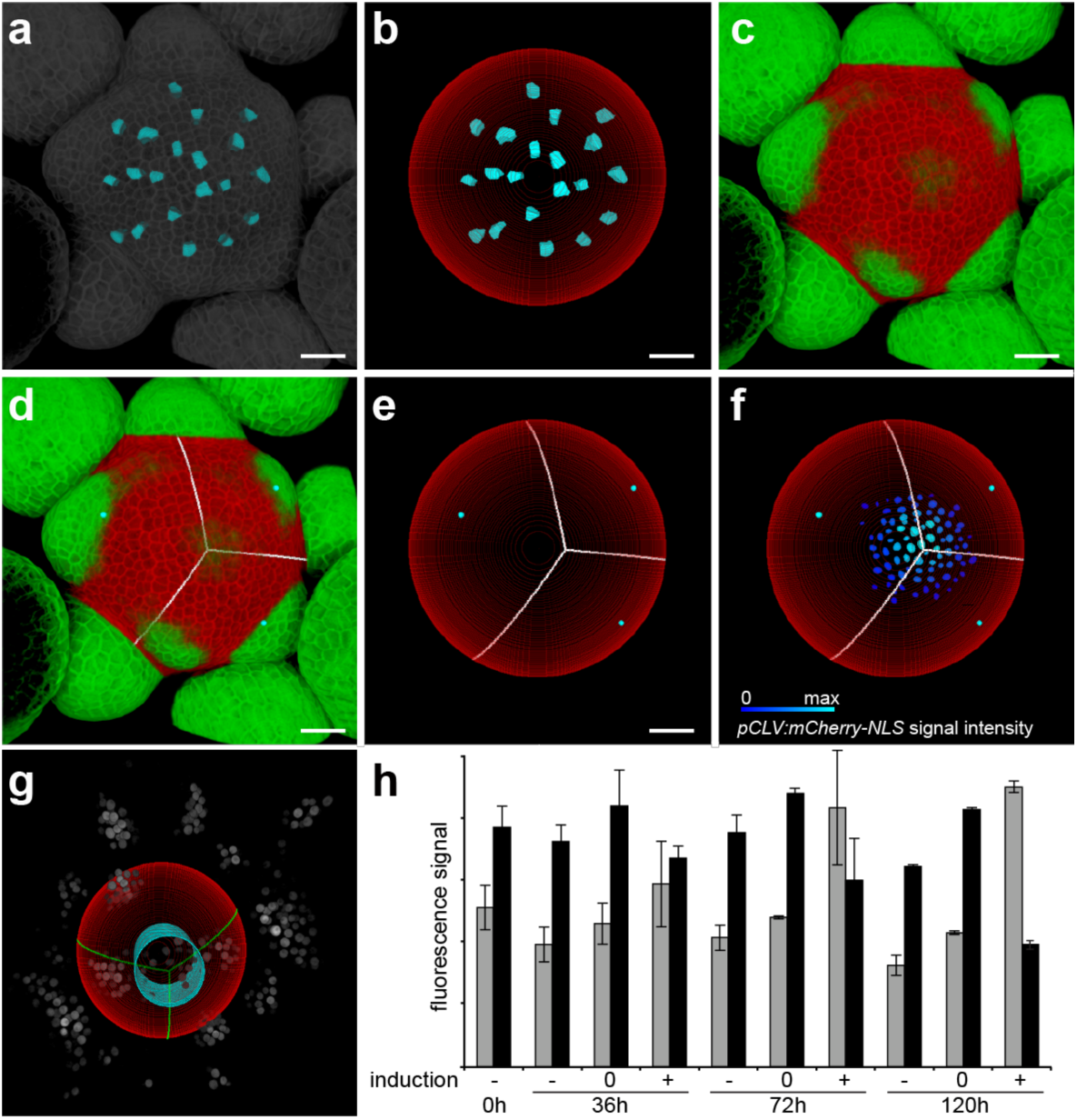
Computational strategy to identify stem cells and DR5v2 quantification. **a)** In a first step, cells across the L1 of the SAM are segmented. **b)** Based on the position of segmented cells, a perfect sphere is fitted to the SAM. **c)** The sphere is applied to the SAM and organ primordia are identified by emergence through the sphere. **d, e)** Equidistant points between the primordia are calculated and used to triangulate the center of the SAM. **f)** The triangulated center was benchmarked against SAMs haboring *pCLV3* reporter labelled stem cells (n×9). The triangulation invariantly identified one of the most central *pCLV3* positive cells. See also Methods. **g)** For signal quantification in the stem cell domain, a cylinder with radius r_cyl_ (× 1/3 * r_sphere_) mimicking the average size of the *CLV3* domain was placed into the computationally identified center of the SAM and fluorescence intensities were quantified within this narrowly defined subdomain. DR5v2-NLS signals are shown in grey, SAM sphere derived from segmentation in red, triangulation lines in green and quantification cylinder in cyan. **h)** Quantification of fluorescent signals from all SAMs of the stem cell loss experiment described in Fig. 2. Total fluorescence signal intensities for *pDR5v2:3xVENUS-NLS* and *pRPS5a:NLS-tdTomato* for the inner region (I_cyl_) and for the peripheral region (I_sphere_) were extracted from respective image volumes. I_cyl_ was averaged over all plants for each time-point and condition and normalized to the overall signal (I_cyl_ + I_sphere_). Grey bars: DR5v2:3xVENUS-NLS signal, black bars: pRPS5a:NLS-tdTomato signal. -: mock treated, 0: ethanol induced, but no observable stem cell loss, +: ethanol induced and stem cell loss. Scale bars: 20*µ*m.

**Supplementary Figure 2:**
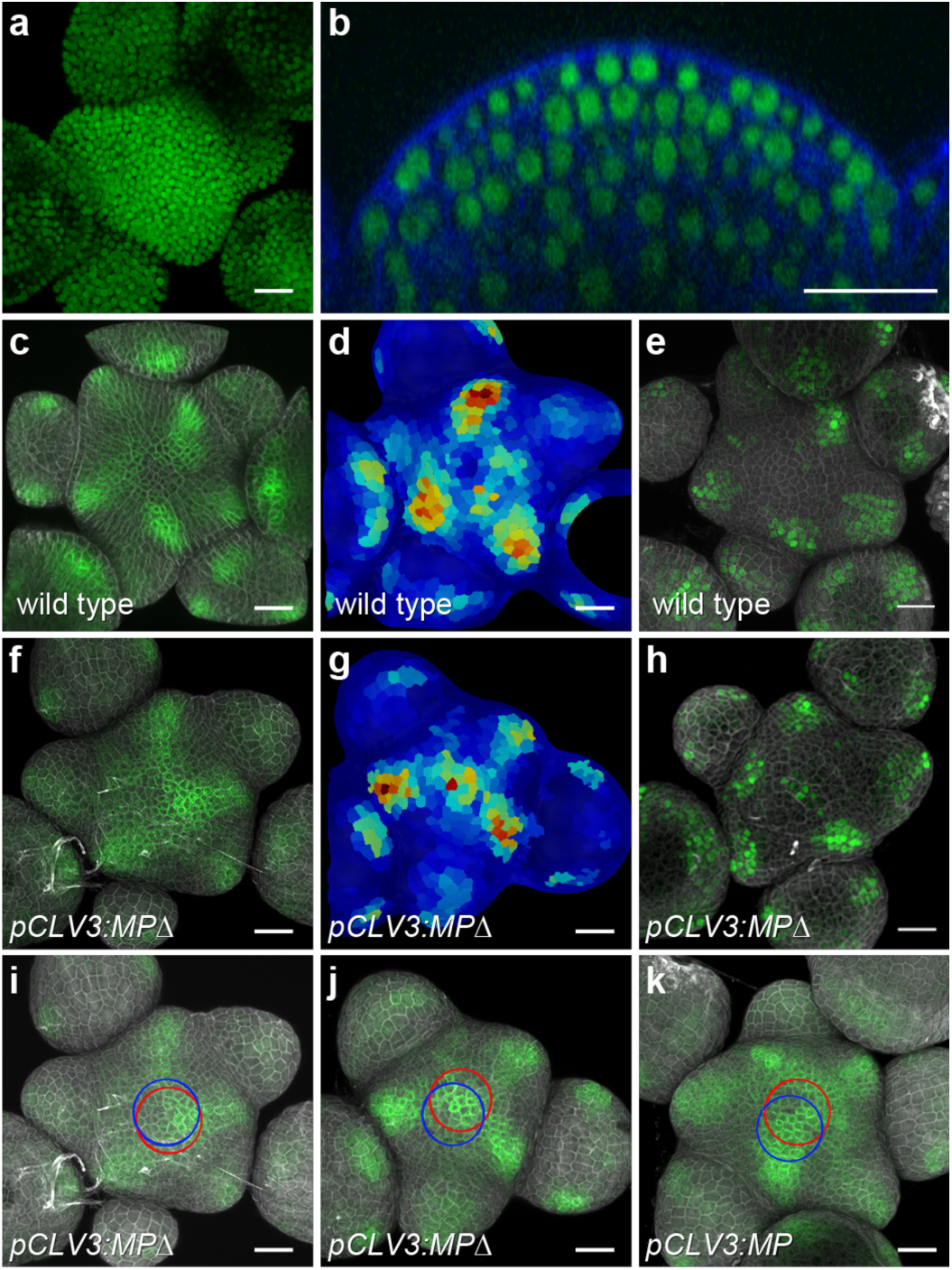
Activity of the *pHMG* promoter, behavior of nuclear and ER localized DR5v2 reporters and auxin signaling output in wild type and *pCLV3:MPΔ* lines. **a, b)** Transgenic line carrying 1347 bp upstream of the At1g76110 locus fused to the *GFP-NLS* coding sequence. **a)** GFP channel in top view. **b)** Side view though a representative SAM showing DAPI and GFP channel. **c)** *pDR5v2:ER-EYFP-HDEL* in wild type. **d)** Per cell quantification of an independent *pDR5v2:ER-EYFP-HDEL* wild-type SAM. **e)** *pDR5v2:3xVENUS-NLS* in wild type. **f-k)** Auxin signaling output was present in the centre of *pCLV3:MP* and *pCLV3:MPΔ* lines, indicated by two independent reporters *pDR5v2:ER-EYFP-HDEL* (6 out of 8 independent T1 plants) (F) and *pDR5v2:3xVENUS-NLS* (6 out of 7 independent T1 plants) (H). **g)** Per cell quantification of *pDR5v2:ER-EYFP-HDEL* in an independent *pCLV3:MPΔ* SAM. DR5v2 activity was not observed in the center of wild-type SAMs grown in the same experiments. **i-k)** Computationally derived central zone in L1 (red) and L3 (blue) are superimposed to SAMs of *pDR5v2:ER-EYFP-HDEL* carrying *pCLV3:MPΔ* (I, J) and *pCLV3:MP* (K). DR5v2 signal clearly coincides with central zone. Scale bars: 20*µ*m.

**Supplementary Figure 3:**
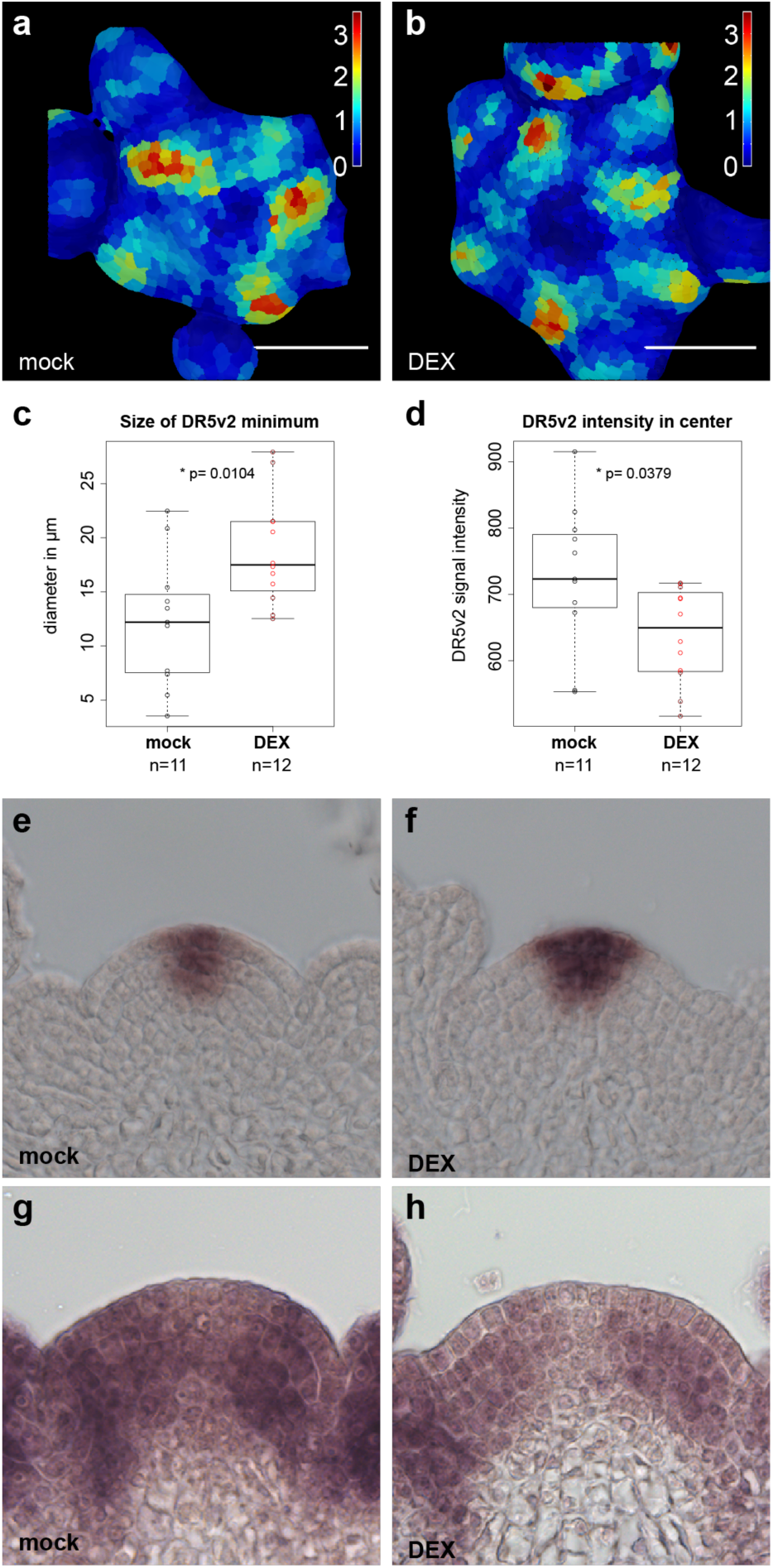
SAM specific molecular responses to ectopic WUS induction. 24 hours after induction of ectopic WUS-GR activity, DR5v2 signal in the central zone was supressed and *CLV3* mRNA expression was enhanced. Representative in situ quantifications of DR5v2 signal after mock **(a)** and DEX **(b)** treatments. **c)** Quantification of the size of the central DR5v2 minimum. **d)** Quantification of the average DR5v2 signal intensity in the central zone. **e)** *CLV3* mRNA expression after 24 hours of mock treatment. **f)** *CLV3* mRNA expression after 24 hours of DEX treatment. **g)** *TIR1* mRNA expression after 24 hours of mock treatment. **h)** *TIR1* mRNA expression after 24 hours of DEX treatment. SAMs of both treatment types were hybridized on the same microscopic slide and imaged under identical settings.

**Supplementary Figure 4:**
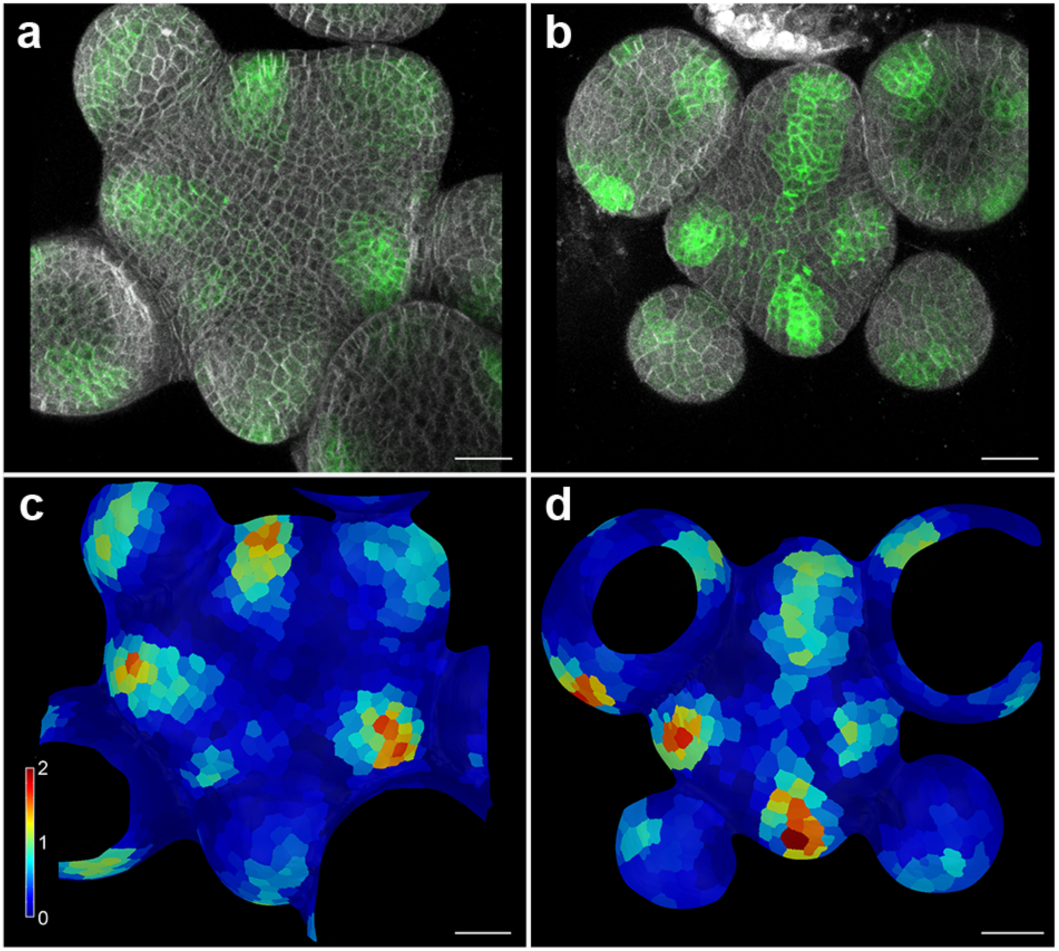
SAMs of *wus-7* plants show auxin signaling output in the stem cell domain. **a)** Representative image of *pDR5v2:ER-eYFP-HDEL* signal in the SAM of L*er* wild-type plants. Only 16% of plants showed DR5v2 activity in the center of the SAM (n×38). **b)** Representative image of *pDR5v2:ER-eYFP-HDEL* signal in a *wus-7* SAM before termination. 61% of *wus-7* plants showed DR5v2 activity in the center of the SAM (n×13). Per cell quantification of DR5v2 signal in wild type (**c**) and *wus-7* (**d**). Scale bars: 20 *µ*m

**Supplementary Figure 5:**
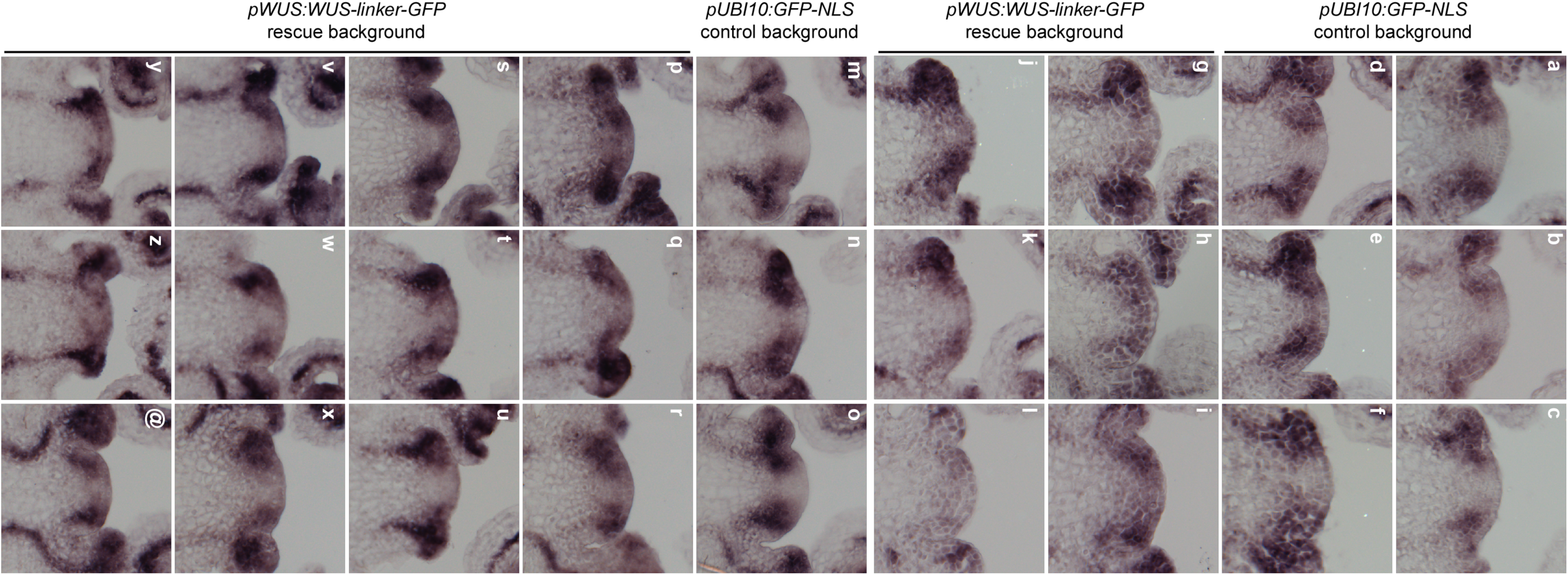
*MP* mRNA expression after induced WUS loss of function. In two independent experiments. **a-l) Experiment I. a-f)** In situ detection of *MP* mRNA in *pUBI10:GFP-NLS* control plants carrying *pCLV3:AlcR/AlcA:NSlmb-vhhGFP4* after 24h of ethanol treatment. **g-l)** In situ detection of *MP* mRNA in stable *pWUS:WUS-linker-GFP wus* rescue plants carrying *pCLV3:AlcR/AlcA:NSlmb-vhhGFP4* after 24h of ethanol treatment. **m-@) Experiment II. m-o)** In situ detection of *MP* mRNA in *pUBI10:GFP-NLS* control plants carrying *pCLV3:AlcR/AlcA:NSlmb-vhhGFP4* after 24h of ethanol treatment. **p-@)** In situ detection of *MP* mRNA in stable *pWUS:WUS-linker-GFP wus* rescue plants carrying *pCLV3:AlcR/AlcA:NSlmb-vhhGFP4* after 24h of ethanol treatment. SAMs of both genotypes were hybridized in sets of two independent experiments and imaged under identical settings. Unadjusted images are shown.

### Supplementary Tables 1-2

**Supplementary Table 1:** GO category enrichment analysis of direct WUS targets. Top 100 enriched categories are shown.

|  | GO ID | Term | Annotated | Significant | Expected | p-Value |
| --- | --- | --- | --- | --- | --- | --- |
| 1 | GO:0010200 | response to chitin | 393 | 145 | 55.45 | 2.8E-30 |
| 2 | GO:0009611 | response to wounding | 313 | 109 | 44.16 | 1E-20 |
| 3 | GO:0010363 | regulation of plant-type hypersensitive response | 336 | 111 | 47.41 | 4.6E-19 |
| 4 | GO:0006612 | protein targeting to membrane | 340 | 111 | 47.97 | 1.3E-18 |
| 5 | GO:0009414 | response to water deprivation | 374 | 130 | 52.77 | 5.7E-18 |
| 6 | GO:0009867 | jasmonic acid mediated signaling pathway | 256 | 89 | 36.12 | 1.2E-15 |
| 7 | GO:0009733 | response to auxin | 354 | 107 | 49.95 | 2.3E-15 |
| 8 | GO:0002679 | respiratory burst involved in defense response | 114 | 50 | 16.09 | 1.1E-14 |
| 9 | GO:0009737 | response to abscisic acid | 548 | 174 | 77.32 | 1.1E-14 |
| 10 | GO:0009738 | abscisic acid-activated signaling pathway | 232 | 78 | 32.74 | 1.1E-12 |
| 11 | GO:0009651 | response to salt stress | 704 | 187 | 99.33 | 2.6E-12 |
| 12 | GO:0009695 | jasmonic acid biosynthetic process | 125 | 49 | 17.64 | 3.4E-12 |
| 13 | GO:0006857 | oligopeptide transport | 97 | 41 | 13.69 | 1.1E-11 |
| 14 | GO:0050832 | defense response to fungus | 303 | 84 | 42.75 | 3.3E-10 |
| 15 | GO:0009862 | systemic acquired resistance, salicylic acid mediated signaling pathway | 222 | 66 | 31.32 | 1.2E-9 |
| 16 | GO:0042538 | hyperosmotic salinity response | 152 | 50 | 21.45 | 2.9E-9 |
| 17 | GO:0009612 | response to mechanical stimulus | 59 | 27 | 8.32 | 4.5E-9 |
| 18 | GO:0042742 | defense response to bacterium | 344 | 93 | 48.54 | 4.9E-9 |
| 19 | GO:0009684 | indoleacetic acid biosynthetic process | 94 | 36 | 13.26 | 5.2E-9 |
| 20 | GO:0006569 | tryptophan catabolic process | 67 | 29 | 9.45 | 6E-9 |
| 21 | GO:0009723 | response to ethylene | 325 | 101 | 45.86 | 1.2E-8 |
| 22 | GO:0009753 | response to jasmonic acid | 427 | 141 | 60.25 | 1.2E-8 |
| 23 | GO:0009873 | ethylene-activated signaling pathway | 118 | 41 | 16.65 | 1.3E-8 |
| 24 | GO:0009620 | response to fungus | 440 | 132 | 62.08 | 2.5E-8 |
| 25 | GO:0000165 | MAPK cascade | 197 | 57 | 27.8 | 4.5E-8 |
| 26 | GO:0009963 | positive regulation of flavonoid biosynthetic process | 93 | 34 | 13.12 | 5.3E-8 |
| 27 | GO:0006355 | regulation of transcription, DNA-templated | 1588 | 296 | 224.07 | 7.1E-8 |
| 28 | GO:0043069 | negative regulation of programmed cell death | 158 | 48 | 22.29 | 1E-7 |
| 29 | GO:0009739 | response to gibberellin | 143 | 49 | 20.18 | 1.1E-7 |
| 30 | GO:0031348 | negative regulation of defense response | 246 | 65 | 34.71 | 2.3E-7 |
| 31 | GO:0009409 | response to cold | 539 | 118 | 76.05 | 4.2E-7 |
| 32 | GO:0009750 | response to fructose | 127 | 39 | 17.92 | 0.0000011 |
| 33 | GO:0030968 | endoplasmic reticulum unfolded protein response | 171 | 48 | 24.13 | 0.0000013 |
| 34 | GO:0009693 | ethylene biosynthetic process | 110 | 35 | 15.52 | 0.0000016 |
| 35 | GO:0009805 | coumarin biosynthetic process | 51 | 21 | 7.2 | 0.000002 |
| 36 | GO:0010310 | regulation of hydrogen peroxide metabolic process | 159 | 45 | 22.43 | 0.0000022 |
| 37 | GO:0030003 | cellular cation homeostasis | 146 | 42 | 20.6 | 0.000003 |
| 38 | GO:0007623 | circadian rhythm | 156 | 44 | 22.01 | 0.0000032 |
| 39 | GO:0006833 | water transport | 118 | 36 | 16.65 | 0.0000034 |
| 40 | GO:0009741 | response to brassinosteroid | 102 | 37 | 14.39 | 0.0000036 |
| 41 | GO:0080167 | response to karrikin | 114 | 35 | 16.09 | 0.000004 |
| 42 | GO:0002237 | response to molecule of bacterial origin | 97 | 31 | 13.69 | 0.0000056 |
| 43 | GO:0006979 | response to oxidative stress | 407 | 90 | 57.43 | 0.0000065 |
| 44 | GO:0006813 | potassium ion transport | 35 | 16 | 4.94 | 0.0000066 |
| 45 | GO:0046777 | protein autophosphorylation | 131 | 37 | 18.48 | 0.000018 |
| 46 | GO:0006598 | polyamine catabolic process | 34 | 15 | 4.8 | 0.000022 |
| 47 | GO:0035556 | intracellular signal transduction | 446 | 133 | 62.93 | 0.000023 |
| 48 | GO:0009269 | response to desiccation | 31 | 14 | 4.37 | 0.00003 |
| 49 | GO:0031347 | regulation of defense response | 485 | 146 | 68.43 | 0.00003 |
| 50 | GO:0009825 | multidimensional cell growth | 96 | 29 | 13.55 | 0.000037 |
| 51 | GO:0009697 | salicylic acid biosynthetic process | 181 | 46 | 25.54 | 0.000037 |
| 52 | GO:0019344 | cysteine biosynthetic process | 181 | 46 | 25.54 | 0.000037 |
| 53 | GO:0006970 | response to osmotic stress | 749 | 207 | 105.68 | 0.000041 |
| 54 | GO:0070838 | divalent metal ion transport | 184 | 53 | 25.96 | 0.000069 |
| 55 | GO:0009627 | systemic acquired resistance | 395 | 109 | 55.73 | 0.000077 |
| 56 | GO:0006949 | syncytium formation | 19 | 10 | 2.68 | 0.000083 |
| 57 | GO:0042398 | cellular modified amino acid biosynthetic process | 50 | 18 | 7.06 | 0.000091 |
| 58 | GO:0009751 | response to salicylic acid | 423 | 122 | 59.69 | 0.000098 |
| 59 | GO:0042631 | cellular response to water deprivation | 59 | 20 | 8.32 | 0.0001 |
| 60 | GO:0009965 | leaf morphogenesis | 186 | 49 | 26.24 | 0.00011 |
| 61 | GO:0010583 | response to cyclopentenone | 132 | 35 | 18.63 | 0.00012 |
| 62 | GO:0001666 | response to hypoxia | 74 | 23 | 10.44 | 0.00014 |
| 63 | GO:0007030 | Golgi organization | 160 | 40 | 22.58 | 0.00017 |
| 64 | GO:0016126 | sterol biosynthetic process | 150 | 38 | 21.17 | 0.00018 |
| 65 | GO:0019748 | secondary metabolic process | 527 | 133 | 74.36 | 0.00022 |
| 66 | GO:0006468 | protein phosphorylation | 620 | 157 | 87.48 | 0.00024 |
| 67 | GO:0006995 | cellular response to nitrogen starvation | 21 | 10 | 2.96 | 0.00024 |
| 68 | GO:0009863 | salicylic acid mediated signaling pathway | 315 | 92 | 44.45 | 0.00028 |
| 69 | GO:0009407 | toxin catabolic process | 180 | 43 | 25.4 | 0.00029 |
| 70 | GO:0009595 | detection of biotic stimulus | 92 | 26 | 12.98 | 0.0003 |
| 71 | GO:0046686 | response to cadmium ion | 415 | 84 | 58.56 | 0.00033 |
| 72 | GO:0006816 | calcium ion transport | 108 | 29 | 15.24 | 0.00036 |
| 73 | GO:0042335 | cuticle development | 42 | 15 | 5.93 | 0.00038 |
| 74 | GO:0009617 | response to bacterium | 499 | 140 | 70.41 | 0.0004 |
| 75 | GO:0010264 | myo-inositol hexakisphosphate biosynthetic process | 51 | 17 | 7.2 | 0.00041 |
| 76 | GO:0010119 | regulation of stomatal movement | 47 | 16 | 6.63 | 0.00046 |
| 77 | GO:0043900 | regulation of multi-organism process | 115 | 30 | 16.23 | 0.00049 |
| 78 | GO:0010017 | red or far-red light signaling pathway | 39 | 14 | 5.5 | 0.00056 |
| 79 | GO:0010260 | animal organ senescence | 27 | 11 | 3.81 | 0.00063 |
| 80 | GO:0009740 | gibberellic acid mediated signaling pathway | 72 | 21 | 10.16 | 0.0007 |
| 81 | GO:0007169 | transmembrane receptor protein tyrosine kinase signaling pathway | 113 | 29 | 15.94 | 0.0008 |
| 82 | GO:0015824 | proline transport | 68 | 20 | 9.59 | 0.00083 |
| 83 | GO:0010227 | floral organ abscission | 32 | 12 | 4.52 | 0.00088 |
| 84 | GO:0052541 | plant-type cell wall<br>cellulose metabolic process | 24 | 10 | 3.39 | 0.0009 |
| 85 | GO:0010158 | abaxial cell fate<br>specification | 7 | 5 | 0.99 | 0.00091 |
| 86 | GO:0009742 | brassinosteroid mediated<br>signaling pathway | 37 | 13 | 5.22 | 0.0011 |
| 87 | GO:0048767 | root hair elongation | 164 | 38 | 23.14 | 0.00117 |
| 88 | GO:0010118 | stomatal movement | 86 | 32 | 12.13 | 0.00168 |
| 89 | GO:0009694 | jasmonic acid metabolic<br>process | 147 | 58 | 20.74 | 0.00171 |
| 90 | GO:0033500 | carbohydrate homeostasis | 12 | 8 | 1.69 | 0.00174 |
| 91 | GO:0007231 | osmosensory signaling<br>pathway | 5 | 4 | 0.71 | 0.00175 |
| 92 | GO:2000022 | regulation of jasmonic acid<br>mediated signaling<br>pathway | 5 | 4 | 0.71 | 0.00175 |
| 93 | GO:0010037 | response to carbon dioxide | 5 | 4 | 0.71 | 0.00175 |
| 94 | GO:0009624 | response to nematode | 72 | 20 | 10.16 | 0.0018 |
| 95 | GO:0006766 | vitamin metabolic process | 77 | 21 | 10.86 | 0.0018 |
| 96 | GO:0006865 | amino acid transport | 228 | 61 | 32.17 | 0.00209 |
| 97 | GO:0000038 | very long-chain fatty acid<br>metabolic process | 44 | 14 | 6.21 | 0.00214 |
| 98 | GO:0046885 | regulation of hormone<br>biosynthetic process | 8 | 5 | 1.13 | 0.00215 |
| 99 | GO:0050801 | ion homeostasis | 205 | 59 | 28.93 | 0.00226 |
| 100 | GO:0052546 | cell wall pectin metabolic<br>process | 40 | 13 | 5.64 | 0.00247 |

**Supplementary Table 2:** Response of genes with activities in auxin signalling to WUS. Adjusted p-value for RNA-seq data was calculated using the Benjamini-Hochberg method in Deseq2. Asterisks denote genes in regions with closed chromatin^26^.

| AGI | Gene Name | # WUS peaks | Log2FC | p adj. |
| --- | --- | --- | --- | --- |
| AT1G59750 | ARF1 | 0 | -0,092540577 | 0,425729231 |
| AT5G62000 | ARF2 | 1 | -0,141313595 | 0,070074796 |
| AT2G33860 | ARF3 | 1 | 0,775921196 | 2,11E-06 |
| AT5G60450 | ARF4 | 1 | -1,258170789 | 4,76E-18 |
| AT1G19850 | ARF5 | 1 | 0,864357040 | 0,000125261 |
| AT1G30330 | ARF6 | 5 | 0,294727963 | 0,014033872 |
| AT5G20730 | ARF7 | 0 | 0,171498773 | 0,24584083 |
| AT5G37020 | ARF8 | 1 | -1,575709325 | 8,52E-20 |
| AT4G23980 | ARF9 | 1 | 1,153767834 | 5,92E-25 |
| AT2G28350 | ARF10 | 2 | 0,847838948 | 0,000948755 |
| AT2G46530 | ARF11 | 1 | 0,925587920 | 4,27E-07 |
| AT1G34310* | ARF12 | 0 | 0 | 1 |
| AT1G34170* | ARF13 | 0 | 0 | 1 |
| AT1G35540* | ARF14 | 0 | 0 | 1 |
| AT1G35520* | ARF15 | 0 | 0 | 1 |
| AT4G30080 | ARF16 | 0 | 0,143765542 | 0,628545091 |
| AT1G77850 | ARF17 | 0 | 0,867761468 | 4,19E-05 |
| AT3G61830* | ARF18 | 0 | 0,989685885 | 7,48E-15 |
| AT1G19220 | ARF19 | 0 | 1,115504426 | 2,96E-09 |
| AT1G35240* | ARF20 | 0 | 0 | 1 |
| AT1G34410* | ARF21 | 0 | 0 | 1 |
| AT1G34390* | ARF22 | 0 | 0 | 1 |
| AT1G43950* | ARF23 | 0 | 0 | 1 |
| AT4G14560 | IAA1 | 1 | -0,026815594 | 0,947757473 |
| AT3G23030 | IAA2 | 2 | -0,763296850 | 2,39E-21 |
| AT1G04240 | SHY2 | 2 | 3,215535318 | 4,66E-121 |
| AT5G43700 | ATAUX2-11 | 1 | -0,449766274 | 2,15E-05 |
| AT1G15580* | IAA5 | 0 | 0 | 1 |
| AT1G52830 | IAA6 | 2 | 0 | 1 |
| AT3G23050 | IAA7 | 1 | 0,572994647 | 4,02E-07 |
| AT2G22670 | IAA8 | 2 | 1,394465988 | 6,40E-43 |
| AT5G65670 | IAA9 | 2 | 0,124793826 | 1,41E-01 |
| AT1G04100* | IAA10 | 0 | -1,709850381 | 2,57E-14 |
| AT4G28640 | IAA11 | 0 | -0,679861940 | 1,26E-02 |
| AT1G04550 | IAA12 | 1 | -0,559301573 | 0,012657873 |
| AT2G33310 | IAA13 | 1 | -0,996497622 | 2,41E-15 |
| AT4G14550 | IAA14 | 2 | -0,578151620 | 0,006372765 |
| AT1G80390 | IAA15 | 0 | 0 | 1 |
| AT3G04730 | IAA16 | 1 | -0,387494366 | 2,49E-09 |
| AT1G04250 | AXR3 | 1 | 0,668789409 | 1,70E-04 |
| AT1G51950 | IAA18 | 3 | -0,752605675 | 2,02E-11 |
| AT3G15540 | IAA19 | 2 | 1,441217799 | 1,10E-03 |
| AT2G46990 | IAA20 | 1 | 1,931969488 | 4,30E-16 |
| AT3G16500 | PAP1 | 2 | -1,484257914 | 1,32E-35 |
| AT4G29080 | PAP2 | 1 | 0,879788470 | 7,44E-07 |
| AT5G25890 | IAA28 | 0 | -0,277043301 | 0,253529454 |
| AT4G32280 | IAA29 | 0 | 1,973159875 | 3,36E-07 |
| AT3G62100 | IAA30 | 0 | 1,183375731 | 9,73E-04 |
| AT3G17600* | IAA31 | 0 | 0 | 1 |
| AT2G01200* | IAA32 | 0 | 2,883104933 | 0,081310628 |
| AT1G15050* | IAA34 | 0 | -0,443724915 | 0,434574757 |
| AT4G03190 | AFB1 | 0 | -1,546414697 | 3,32E-12 |
| AT3G26810 | AFB2 | 2 | -0,145179846 | 3,61E-01 |
| AT1G12820 | AFB3 | 0 | 1,196545030 | 1,29E-40 |
| AT4G24390 | AFB4 | 0 | 0,581458460 | 0,001034196 |
| AT5G49980 | AFB5 | 1 | 0,477719550 | 4,95E-05 |
| AT3G62980 | TIR1 | 1 | -0,871245173 | 4,91E-18 |
| AT1G73590 | PIN1 | 1 | 0,133975925 | 0,771092197 |
| AT5G57090 | PIN2 | 0 | 0,678755084 | 0,07343843 |
| AT1G70940 | PIN3 | 2 | -1,171670670 | 1,85E-27 |
| AT2G01420 | PIN4 | 3 | 0,364037027 | 2,92E-05 |
| AT5G16530* | PIN5 | 0 | -0,976327009 | 0,792123994 |
| AT1G77110 | PIN6 | 1 | 1,567558864 | 0,056844235 |
| AT1G23080 | PIN7 | 2 | -0,170157356 | 0,304697713 |
| AT5G15100 | PIN8 | 0 | 0 | 1 |
| AT2G38120 | AUX1 | 2 | 0,905501335 | 3,36E-10 |
| AT5G01240 | LAX1 | 2 | 0,159143298 | 0,207550001 |
| AT2G21050 | LAX2 | 0 | 0,461205197 | 0,145235679 |
| AT1G77690 | LAX3 | 1 | 0,011696012 | 0,965073056 |
| AT2G34650 | PID | 2 | 0,345593382 | 0,104333098 |
| AT2G26700 | PID2 | 0 | -0,114850869 | 0,830691073 |

## Materials and Methods

### Plant material and treatments

All plants were grown at 23 °C in long days or continuous light. Ethanol inductions were performed by watering with 1% ethanol and continuous exposure to ethanol vapour, refreshed every 12 hours. WUS-GR was induced by submerging seedlings in 25 μM dexamethasone, 0.015% Silwet L-70 in 0.5x MS for 2 hours. For local induction at the SAM, 10 *µ*l induction solution were directly applied to the primary inflorescence meristem. Auxin plates were 0.5x MS, 1% agar, pH 5.7, 10 *µ*m IAA. For TSA/IAA cotreatments, shoot apical meristems were dissected from about 4 cm high stem and cultured *in vitro* in Apex Growth Medium (AGM) overnight^42^. AGM was supplemented with vitamins (Duchefa M0409), cytokinin (200 nM 6-Benzylaminopurine), and IAA (3-indole acetic acid, 1 mM) and/or Trichostatin A (TSA, Sigma, T8552, final concentration 5 *µ*M) or mock before pouring. IAA stock solution (0.1 M in 0.2 M KOH) was diluted with 2 mM M.E.S (pH 5.8) to 1 mM working solution, then added to the plates for 30 min before imaging on the second day.

For WUS-induction with TSA treatments, seedlings were submerged in DEX (10 *µ*M) or TSA (1 *µ*M) solution or both, slowly shaken for 2 h, and then harvested for RNA-seq.

All plants were of Col-0 accession apart from *wus-7*, which was in L*er* background. For experiments involving *wus-7*, L*er* plants were used as controls.

### Transgenes

The *R2D2* and *pDR5v2:3xVENUS-NLS* lines have been described in ref. 16. *pDR5v2:tdTomato-Linker-NLS:trbcS* was transformed into heterozygous *wus-7* plants and L*er* control plants and activity patterns were scored in T1. A stable single insertion T3 line of *pDR5v2:ER-EYFP-HDEL:tAt4g24550* was used for transformation with *pCLV3:3xmCherry-NLS* and signals were scored in T1. For deGradFP the anti-GFP nanobody coding sequence (*NSlmb-vhhGFP4)*^22^ was brought under control of the AlcR/AlcA system^43^ and transformed into a stable *pWUS:WUS-linker-GFP wus* rescue line (GD44, described in ref. 5), or an *pUBI10:GFP-NLS* line as control. Experiments were performed in stable single insertion T3 lines. Similarly, the *pCLV3:AlcR/AlcA:CalS3m* line^5^ was crossed to *pDR5v2:3xVENUS-NLS, pRPS5a:NLS-tdTomato* and F3 single insertion progeny was used for experiments. For ectopic WUS induction lines *mCherry* was fused N-terminally to the ligand-binding domain of the rat glucocorticoid receptor (GR) and linked by (AAASAIAS[SG]11SAAA) to the *WUS* coding sequence under control of the *pUBI10* promoter. A single insertion homozygous line was used for crossings, in RNA-seq, and ChIP-seq.

The *pHMG* promoter corresponds to 1347 bp upstream of the AT1g76110 locus. Most constructs were assembled using GreenGate cloning^44^.

### Microscopy

Confocal microscopy was carried out on a Nikon A1 Confocal with a CFI Apo LWD 25× water immersion objective (Nikon Instruments) as described in ref. 5. 1 mg/ml DAPI was used for cell wall staining.

### Image analysis

Quantitative image analysis was done on isotropic image stacks using Fiji (v1.50b) ^45^, MorphoGraphX^46^, ilastik^47^, Matlab (Release 2014b, The MathWorks, Inc., United States) and KNIME^48^. Signal quantification methods: all images for an experimental set were captured under identical microscope settings and signal intensities were never adjusted, making intra-experiment signal comparisons possible. MorphographX analysis was performed according to standards defined in the user manual. Averaging and statistical analysis of signals across meristems was performed as follows: histograms of signal intensities along 100 central cross-sections per SAM were (cross-sections rotated by 3.6 degrees successively) were measured by ImageJ standard function. Signals were centered for comparison between individuals. Signals +/-12.5*µ*m around the SAM center were compared between treatment and control and tested for significance by Student’s T-test. Distance from center with signal up to 120% of center background signal between treatment and control was determined and tested by Student’s T-test.

To determine the center of an inflorescence meristem, 10 to 20 L1 cells located at the meristem summit were segmented using the carving workflow in ilastik. A sphere was fitted through the centroids of these cells using the least squared distances method. The sphere was superimposed on the original DAPI stained image volume to help identifying the newly emerging flower primordia. Three points marking the center of three young flower primordia were manually picked close to the sphere surface, projected onto the sphere and then used as seeds to perform a spheric voronoi tessellation (https://de.mathworks.com/matlabcentral/fileexchange/40989-voronoi-sphere). The point P_center_ is equidistant to the three seed points and serves as a good approximation for the meristem center which is marked by the *pCLV3* stem cell reporter. The method was tested using image stacks of nine meristems containing cell walls stained by DAPI in one channel and the stem cell marker *pCLV3::mCherry-NLS* in the second channel. The computationally estimated meristem center and the one determined by *pCLV3:mCherry-NLS* expression in every case were in the range of one cell diameter. Further details and workflows are available on request.

### In situ hybridization

In-situ hybridizations were carried out as described in ref. 49.

### ChIP-seq and RNA-seq

All experiments were carried out on 5 day old seedlings grown on 0.5 MS plates after 2 hours of either Dex or mock treatment. ChIP assays were performed from 3g of fresh weight each as described in ref. 50 using RFP-Trap single chain antibodies (Chromotek). Enrichment of specific DNA fragments was validated by qPCR at the *ARR7* promoter region^24^. Two independent libraries were generated for the *WUS-GR* and control ChIP each using pooled DNA from 6 to 9 individual ChIP preparations. RNA-seq was carried out in biological triplicates. After careful benchmarking of our WUS-GR line, we find it to be the most potent and consistent tool for WUS induction to date, affording a much higher sensitivity for identifying transcriptional targets. In addition, the use of RFP-trap increased sensitivity of the ChIP assay. Consistently, we were able to identify 6740 genomic regions bound by WUS in both ChIP-seq experiments at p< 0.05. This compared to 136 regions we had previously identified by ChIP-chip^23^, highlighting the increase in power. Previously identified direct targets, such as *ARR7, CLV1, KAN1, KAN2 AS2* and *YAB3*^23-25^ were also picked up in our analysis. Because of the medium level ubiquitous expression of WUS, both RNA-seq and ChIP-seq capture the global regulatory potential of WUS. Since regulatory output of WUS is dependent on tissue context, targets identified here might not be relevant for all tissues. In addition, targets might be induced by WUS in one tissue and repressed in another, which cannot be resolved by this dataset. All genomic datasets are available under GEO accession: GSE122611

### Bioinformatics

ChIP-seq data were mapped to TAIR10 genome by BWA aligner (v0.7.17)^51^ on a local Galaxy instance (v17.09)^52^. Peak calling was performed using Hiddendomains (v3.0)^53^. Peaks were annotated to TAIR10 genes using PAVIS^54^.

Alignment of RNA-seq reads to TAIR10 genome by HISAT2 (v2.1.0)^55^ and calculation of count matrices by featureCounts (v1.6.3)^56^ was done on Galaxy instance. Differentially expressed genes were identified with R bioconductor package Deseq2 (1.20.0)^57^. Gene ontology analysis was carried out using topGO R package (v2.32.0) with all genes annotated to open chromatin^26^ as background.

